# Proteomic evidence of depression-associated astrocytic dysfunction in the human male olfactory bulb

**DOI:** 10.1101/2023.10.29.564604

**Authors:** Reza Rahimian, Kelly Perlman, Gohar Fakhfouri, Refilwe Mpai, Vincent R. Richard, Christa Hercher, Lucy Penney, Maria Antonietta Davoli, Corina Nagy, René P. Zahedi, Christoph H. Borchers, Bruno Giros, Gustavo Turecki, Naguib Mechawar

**Author notes:** Corresponding author: Naguib Mechawar, PhD Douglas Mental Health University Institute, 6875 Boulevard LaSalle, Montréal, Quebec Canada, H4H 1R3. These authors contributed equally (Reza Rahimian, Kelly Perlman, Gohar Fakhfouri).

## Abstract

The olfactory bulb (OB), a major structure of the limbic system, has been understudied in human investigations of psychopathologies such as depression. To explore more directly the molecular features of the OB in depression, a global comparative proteome analysis was carried out with human post-mortem OB samples from 11 males having suffered from depression and 12 healthy controls. We identified 188 differentially abundant proteins (with adjusted p<0.05) between depressed cases and controls. Gene ontology and gene enrichment analyses suggested that these proteins are involved in biological processes including the complement and coagulation cascades. Cell type enrichment analysis displayed a significant reduction in several canonical astrocytic proteins in OBs from depressed patients. Furthermore, using RNA-fluorescence *in-situ* hybridization, we observed a decrease in the percentage of ALDH1L1^+^ cells expressing canonical astrocytic markers including *ALDOC*, *NFIA*, *GJA1 (connexin 43)* and *SLC1A3 (EAAT1)*. These results are consistent with previous reports of downregulated astrocytic marker expression in other brain regions in depressed patients. We also conducted a comparative phosphoproteomic analysis of OB samples and found a dysregulation of proteins involved in neuronal and astrocytic functions. To determine whether OB astrocytic abnormalities is specific to humans, we also performed proteomics on the OB of socially defeated male mice, a commonly used model of depression. Cell-type specific analysis revealed that in socially defeated animals, the most striking OB protein alterations were associated with oligodendrocyte-lineage cells rather than with astrocytes, highlighting an important species difference. Overall, this study further highlights cerebral astrocytic abnormalities as a consistent feature of depression in humans.

## INTRODUCTION

The olfactory bulb (OB) is a major component of the limbic system, a group of brain structures involved in motivation, emotion, learning, and memory (Milardi et al., 2017). The OB transmits olfactory information to other limbic areas, including the amygdala, hippocampus, and the anterior cingulate cortex (Milardi et al., 2017). Different brain regions that participate in the processing of olfactory information have been implicated in depression (Croy and Hummel, 2017; Kohli et al., 2016; Rottstaedt et al., 2018). Interestingly, humans with OB aplasia (congenital anosmia) have higher rates of depressive symptoms (Croy et al., 2012; Rottstaedt et al., 2018), and patients with depression have been reported to display decreased olfactory sensitivity and discrimination (Lombion-Pouthier et al., 2006; Negoias et al., 2010). Moreover, structural MRI has revealed a smaller OB volume in depressed patients compared to healthy controls (Negoias et al., 2010). In rodents, bilateral olfactory bulbectomy results in a depression-like phenotype and chronic antidepressant administration can rescue the associated behavioral and physiological changes (Kelly et al., 1997; Song and Leonard, 2005). Given the highly evolutionarily conserved OB-limbic circuitry, and the resulting functional connections between the olfactory, salience, and emotional brain networks (Croy and Hummel, 2017; Kohli et al., 2016; Rottstaedt et al., 2018; Takahashi et al., 2016), these observations strengthen the hypothesis that depression is associated with impaired OB structure and function. The underlying cellular and molecular changes, however, remain largely unexplored in humans. To examine the molecular alterations occurring in the OB in depression, the present study aimed at performing the first unbiased global proteome and phosphoproteome analysis of post-mortem OB samples from depressed individuals and matched healthy controls. For comparative purposes, we the same proteomics analysis experiments in the OB of socially defeated mice. Our main hypothesis was that the protein profile of canonical glial markers would be altered in the OBs of humans having suffered from depression and in the OBs of socially defeated mice.

## MATERIAL AND METHODS

### Human OB samples

Post-mortem OB samples from depressed patients (D) and healthy controls (CTRL) were obtained from the Douglas-Bell Canada Brain Bank (DBCBB; Montreal, Canada). The DBCBB collaborates closely with the Quebec Coroner’s Office, and all phenotypic information was obtained via standardized psychological autopsies, with informed consent provided by the next of kin (Dumais et al., 2005). Either presence or suspected presence of any neurological or neurodegenerative disorders based on clinical files was an exclusion criterion for this study. Information from the Coroner’s office, medical records, social services records, and toxicological analysis were complemented with proxy-based interviews with one or more informants best acquainted with the donor. Clinical vignettes realized by combining all this information were evaluated by a panel of clinicians, using Diagnostic and Statistical Manual of Mental Disorders (DSM-IV) criteria, to establish a diagnosis (Dumais et al., 2005; Tanti et al., 2022).

Three distinct sets of fresh-frozen OB samples from depressed patients and healthy controls were used in this study: the first for proteomic (12 CTRL, 11 D) and phosphoproteomic (analyses of 9 samples/group), the second for RNA-fluorescent *in situ* hybridization (5 CTRL, 3 D), and the third for immunofluorescence (4 CTRL, 5 D). Unfortunately, although the same OB lysate was used for proteomic and phosphoproteomic analyses, phosphoprotein enrichments were only successful in 9 samples per group. Group characteristics, including age, post-mortem interval (PMI), tissue pH, substance dependence, and antidepressant use, are presented in Supplementary Tables 1, 2, 3, and 4.

### Chronic social defeat stress and mouse OB samples

Animal experiments were conducted with the approval of the Douglas Animal Care Committee. Male C57BL/6 mice were subjected to a social defeat paradigm, consisting of a daily 5-minute defeat with a different CD-1 aggressor mouse on each of 10 consecutive days (Golden et al., 2011) (all details are provided in the Supplementary Methods). Twenty-four hours following the last behavioral test, socially defeated and control mice were euthanized, the OB pairs were extracted on an ice-cold metal surface, snap frozen and kept at -80°C for proteomics analysis. For the social defeated group, only susceptible mice consistently displaying susceptibility throughout the behavioral tests (Open Field, Social Interaction and Elevated Plus Maze; all details are provided in the Supplementary Methods) following the chronic stress paradigm were chosen. Every two pairs of OBs were pooled to serve as a single biological replicate.

### Proteomic and phosphoproteomic experiments and analyses

For both human and mouse samples, proteins were extracted from whole frozen OBs in tissue lysis buffer containing 5% SDS (Fisher Chemical), 100 mM TRIS (Fisher BioReagents™) pH 8.5 supplemented with PhosStop phosphatase inhibitor as well as cOmplete™ mini protease inhibitor cocktail (Roche). Samples were subjected to probe-based sonication using a Thermo Sonic Dismembrator and subsequently heated at 95° C for 10 minutes. A portion of the sample (less than 5% by volume) was reserved for total protein quantitation by BCA assay (Thermo/Pierce). Disulfide bonds were reduced and alkylated with 20 mM TCEP and 25 mM iodoacetamide. 50 μg of protein per sample was digested with trypsin (Promega) overnight at 37°C using S-TRAP micro cartridges (Protifi LLC,). Peptide-containing samples were desalted using Waters Oasis HLB SPE cartridges. 10% of the sample was reserved for measurement of the total proteome, and the remainder was used for Fe-(III)-NTA based immobilized metal affinity chromatography (IMAC)-based automated phosphopeptide enrichment using an Agilent Bravo liquid handling robot and Agilent AssayMap IMAC cartridges (Beck et al., 2017; Burkhart et al., 2012; Gonczarowska-Jorge et al., 2017). Samples were analyzed by nano Liquid Chromatography with tandem mass spectrometry (LC-MS/MS) by data dependent acquisition (DDA) using an Easy nLC 1200 coupled with a Q Exactive Plus mass spectrometer (Thermo Fisher Scientific).

Samples were first loaded onto a pre-column (Acclaim PepMap 100 C18, 3 µm particle size, 75 µm inner diameter x 2 cm length) in 0.1% formic acid (buffer A). Peptides were then separated using either a 100-min binary gradient for total ranging from 3-40% B (84% acetonitrile, 0.1% formic acid) on the analytical column (Acclaim PepMap 100 C18, 2 µm particle size, 75 µm inner diameter x 25 cm length) at 300 nL/min. MS spectra were acquired from m/z 350-1,500 at a resolution of 70,000, with an automatic gain control (AGC) target of 1 x 106 ions and a maximum injection time of 50 ms. The 15 most intense ions (charge states +2 to +4) were isolated with a window of m/z 1.2, an AGC target of 2 x 104 and a maximum injection time of 64 ms and fragmented using a normalized higher-energy collisional dissociation (HCD) energy of 28. MS/MS spectra were acquired at a resolution of 17,500 and the dynamic exclusion was set to 30 s. DDA MS raw data was processed with Proteome Discoverer 2.5 (Thermo Scientific) and searched using Sequest HT against a FASTA file containing all reviewed protein sequences of the canonical human proteome without isoforms downloaded from Uniprot. The enzyme specificity was set to trypsin with a maximum of 2 missed cleavages. Carbamidomethylation of cysteine was set as static modification and methionine oxidation as variable modification. The precursor ion mass tolerance was set to 10 ppm, and the product ion mass tolerance was set to 0.02 Da. The percolator node was used, and the data was filtered using a false discovery rate (FDR) cut-off of 1% at both the peptide and protein level. The Minora feature detector node of Proteome Discoverer was used for precursor-based label free quantitation (Beck et al., 2017; Burkhart et al., 2012; Gonczarowska-Jorge et al., 2017). Only peptides that are unique to a protein were used for quantitation. In cases where a protein was not found in a particular sample, we imputed the missing areas using Low Abundance Resampling, an algorithm that randomly uses an area value from the lowest 5% of those detected (Gonczarowska-Jorge et al., 2017).

For human samples, proteins or phosphoprotein modifications that were not detected in at least 50% of subjects in both cases and controls were removed. This conservative approach was chosen to increase the robustness of our results given the small sample size, as it ensured that each protein had absolute quantities that were as representative of the group as possible (i.e., no imputation required) without excluding too many proteins. For mouse samples, we elected to remove any protein that was not detected in at least 1/3 of the samples from both groups, since each sample was derived from the combination of 2 OBs and inter-subject variability was lower (6 OBs in control and 6 OBs in social defeated mice). The underlying data structure was visualized using PCA and Grubb’s test was used to identify outliers for removal. Differentially abundant proteins (DAP), referring to proteins whose abundances are significantly different between cases and controls, were computed with t-tests that have “background” adjusted p-values which means that a weighting factor is applied to the p-values that takes into account the distribution of ratios for all other proteins as well as the protein abundances themselves (Navarro et al., 2014). P-values were corrected for multiple comparisons with the Benjamini-Hochberg method. This background t-test is designed to fulfill the assumption of data normality as it handles data such that the variance is constant for the reference points of the background. All downstream analyses and data visualizations were conducted in R. Gene ontology and gene set enrichment analyses were conducted with the clusterProfiler and DOSE packages (Korotkevich et al., 2021; Yu et al., 2012; Yu et al., 2015). Ridgeplots are used visualize top terms as determined by the gseGO function, which was ran with the parameter values used in the online vignette (https://learn.gencore.bio.nyu.edu/rna-seq-analysis/gene-set-enrichment-analysis/). DAPs were compared with the literature using the DisGeNet database (Piñero et al., 2017) and assessed for overlap between our DAPs and genes that were established to be differentially expressed in the following conditions: Mental Depression (CUI: C0011570), Unipolar Depression (CUI: C0041696), Endogenous depression (CUI: C0011573), Severe depression (CUI: C0588008), Depression and Suicide (CUI: C1524032), clinical depression (CUI: C2362914), Major Depressive Disorder (CUI: C1269683), Major depression (CUI: C1269683), provided that the gda score was greater than 0.01. Over enrichment of cell types based on DAPs was calculated by comparing published sets of gene lists and using fisher’s exact test to look for statistically significant overabundance. The list of genes for each major cell type was formed by taking the union of the markers from the McKenzie et al. (2018) and Qiu et al. (2021), with corresponding MarkerPen tool, plus genes selected to complete the cell type categories which were previously deemed insufficient, primary the oligodendrocytes (OLs).

### RNA-fluorescence in-situ hybridization (FISH) in human OB samples

Frozen OBs were cut in serial 10 µm-thick dorsoventral sections with a cryostat, and sections were collected on Superfrost charged slides. *In situ* hybridization was performed using Advanced Cell Diagnostics RNAscope® probes and reagents following the manufacturer’s instructions (Reemst et al., 2022). Sections were first fixed in cold 10% neutral buffered formalin for 15 min, dehydrated by increasing gradient of ethanol baths, and air-dried for 5 min.

Endogenous peroxidase activity was quenched with hydrogen peroxide for 10 min followed by protease digestion for 30 min at room temperature (Tanti et al., 2022). The following probes were then hybridized for 2 h at 40 °C in a humidity-controlled oven: Hs-ALDH1L1 (ACDbio, REF;438881), Hs-ALDOC (ACDbio, REF;407031), Hs-GJA1(ACDbio, REF;444281), Hs-NFIA (ACDbio, REF;442781) and Hs-SLC1A3 (ACDbio, REF;461081). Amplifiers were added using the AMP reagents and the signal visualized through probe-specific HRP-based detection by tyramide signal amplification (TSA, ACDbio) with Opal dyes (Opal 520 and Opal 570; Perkin Elmer) diluted 1:900. In order to eliminate endogenous autofluorescence from lipofuscin and cellular debris, sections were incubated with TrueBlack (Biotium, Cat; 23007) for 30 seconds. Slides were then cover slipped with Vectashield mounting medium with 4′,6-diamidino-2- phenylindole (DAPI) for nuclear staining (Vector Laboratories, H-1800) and kept at 4 °C until imaging. Both positive and negative controls provided by the supplier (ACDbio) were used on separate sections to confirm signal specificity. The slides were imaged at 20X magnification using the Olympus VS120 virtual slide scanner and the scans were transferred to QuPath (v.0.3.0) for further analysis. OB area (whole section) was demarcated manually and QuPath was employed for automated cell detection based on DAPI staining and RNAscope signal quantification. For each probe, only cells bearing three or more fluorescent puncta were counted as positively labeled.

### Immunofluorescence in human OB samples

OBs were placed in 10% neutral buffered formalin for 2 hours. Samples were then stored in a 30% sucrose solution until they were processed for cryosectioning and flash-frozen in isopentane. Samples were then sectioned at 30 μm on a cryostat, collected on charged Super Frost Plus slides and stored at -80 . Slides were rinsed in phosphate-buffered saline (PBS) and incubated in a blocking solution of PBS, 0.2% Triton-X and 10% normal donkey serum (NDS) for an hour at 37 . Following this, sections were briefly rinsed in PBS before being incubated overnight at 4 with an anti-ALDH1L1 antibody (Millipore, 1:250, MABN495 | Proteintech, 1:500, 17390-1-AP) in a blocking solution of PBS/0.2% Triton-X/5% NDS. Slides were then rinsed and incubated for 1 hour at room temperature with the secondary Alexa-647 anti-mouse antibody (Jackson ImmunoResearch, 1:500, 715-605-151) diluted in the same blocking solution. Next, sections were rinsed and endogenous autofluorescence from lipofuscin and cellular debris were quenched with TrueBlack (Biotium) and coverslipped with Vectashield mounting medium (Vector Laboratories, H-1800). For ALDH1L1 quantification, sections were scanned using a Slide Scanner Olympus VS120 and ALDH1L1^+^ astrocytes then quantified using Qupath (Bankhead, 2017). For each subject, two OB sections per subject were quantified. To minimize variability the dorsal most and ventral most sections from each OB were used. Within each section, two ROIs were selected such that both the right and the left part of each tissue section was sampled. Each ROI comprised of 28 250x250 μm grid squares. Within each ROI, all ALDH1L1^+^ cells were counted in the same z-plane. The presence of a DAPI-labelled nucleus overlapping with the ALDH1L1^+^ immunolabelling was required for positive identification of an astrocyte. Group characteristics, including age, PMI, tissue pH, substance dependence, and medication, are presented in Supplementary Tables 4.

### Statistical analysis

R and GraphPad Prism (v8) were employed to perform statistical analyses. For animal studies and FISH experiments, unpaired Student *t* tests were used to compare different behavioral parameters and quantifications, respectively. For FISH quantifications, double- labeled astrocytes were assessed using QuPath (v0.3.2) and reported as a percentage of total astrocytes (ALDH1L1^+^). *p* < 0.05 was set as the threshold for statistical significance in all analyses.

## RESULTS

### Altered expression of canonical astrocyte proteins in OB of depressed individuals

We identified 188 DAPs (with corrected p<0.05 for group differences) between cases and controls, which was narrowed down to 115 proteins when using a 2-sigma fold change cut-off (Figure 1A).

**Figure 1.**
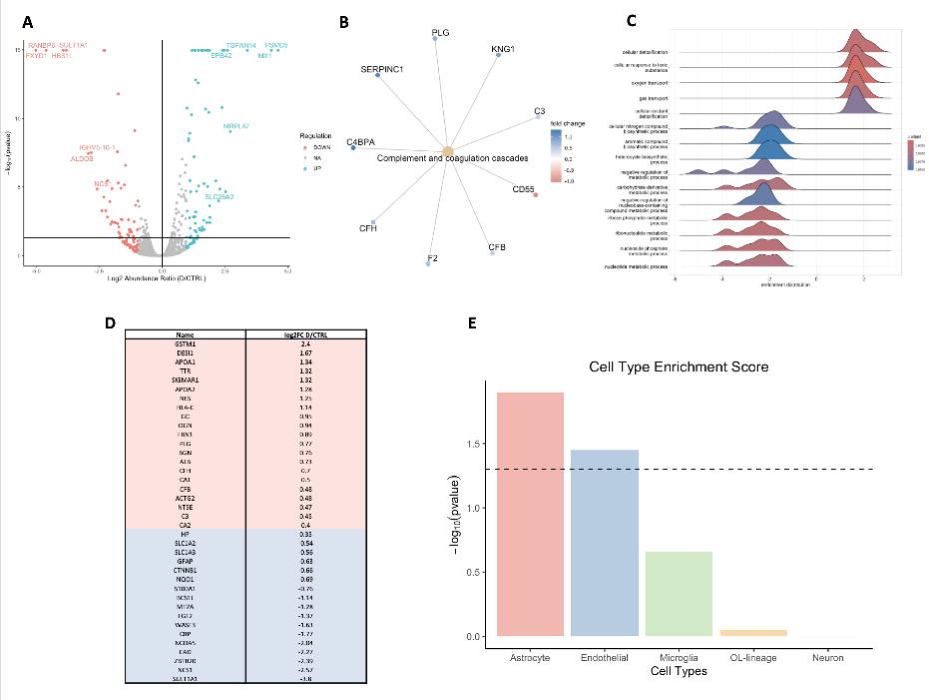
Human OB proteomics reveal enrichment in astrocyte-specific differentially abundant proteins in depressed individuals. (**A**) Volcano plot with log base 2 abundance ratio between cases and controls on the x-axis and the -log_10_(p-value) on the y-axis. Each dot represents a protein, and they are colored by direction of regulation with downregulation in pink and upregulation in turquoise. The log base 2 abundance ratio cutoff is |2.5| for visualization purposes and the p-value is cutoff is -log_10_(0.05). D = Depression, CTRL = Control. (**B**) Graphical representation of the DAPs contributing to the enrichment of the complement and coagulation cascade biological process identified by the enrichKEGG function. The node color of the DAP represents the log2 fold change. (**C**) Ridge plot with the enrichment of the gene ontology (GO) terms on the x-axis, and each GO biological process listed on the y-axis. This plot shows the over enriched GO terms in the list of differentially abundant proteins between cases and controls. The ridges are colored by adjusted p-value of the enrichment tests as demonstrated by the color bar. The positive enrichment distribution values indicate enrichment of the GO term in cases compared to controls, and the negative enrichment distribution values indicate enrichment in controls compared to cases. (**D**) Table of overlap of proteins in list of differentially abundant proteins and genes found in the DisGeNET database. Proteins are listed in descending order by log base 2-fold change between cases and controls. Proteins with positive fold change values are colored in pink and negative fold change values are colored in blue. (**E**) Bar plot showing cell type enrichment scores of differentially abundant proteins. The x-axis shows the broad cell types of the brain, and the y-axis shows the -log_10_(p-value), where the p-value refers to the enrichment p-value from hypergeometric tests. The dotted line refers to the significance cutoff which is -log_10_(0.05). Only the astrocytes surpass this significant threshold.

Supplementary Table 5 shows the number of proteins detected before and after the selected conservative cut-off described in the methods. Gene set enrichment analyses suggested that the DAPs are involved in general biological processes such as cellular detoxification (e.g., GSTM1, HBG1, HBD) and nucleotide metabolic process (e.g., CMPK2, CDAKD, ALDOB, SULT1A1) but the KEGG enrichment tool suggested specific enrichment for complement and coagulation cascade (e.g., C3, F2, PLG, C4BPA, CFH, CFB). Given the scarcity of molecular OB research in depression, we sought to assess the extent to which the list of DAPs overlapped with differentially expressed genes reported in the literature for other brain regions in humans. To this end, we used the DisGeNeT database and considered the differentially expressed genes that were common to studies of depressive disorders. We then interrogated the overlap (i.e., intersection) of those genes with the DAPs identified in the current study and found 38 overlapping candidates from our initial total list of 188 proteins (Figure 1D). These 38 candidates included genes involved in the complement cascade, lipoprotein regulation, and response to oxidative stress. Since the proteomics analyses were performed on bulk OB tissue, we examined whether the DAP list would suggest any cell type specificity to the alterations. We assessed the overlap between broad cell type specific marker genes (astrocyte, endothelial, microglial, OL-lineage, and neuronal) provided in Supplementary Table 6, constructed as described in the Methods section. We applied hypergeometric tests on this list to determine cell type enrichment and found that the only significantly enriched cell types were the astrocyte and endothelial cells (astrocyte, *p* = 0.0128; endothelial, *p* = 0.0356; microglia, *p* = 0.220; OL-lineage, *p* = 0.893; neuron, *p* = 0.993; Figure 1E). However, since the PMI was not matched between groups (Supplementary Table 2), we elected to validate our findings with an alternative analysis. We ran linear models with the following structure: Protein ∼ Group + Age + PMI. We selected proteins whose models had p<0.05 for the group factor as the DAPs. We then conducted the cell type marker enrichment analysis with these new DAPs and found that only astrocyte markers reached significance (astrocyte, *p* = 0.0249; endothelial, *p* = 0.896; microglia, *p* = 0.118; OL-lineage, *p* = 0.986; neuron, *p* = 0.495) (Supplementary Figure 1). Given that astrocyte markers were found to be significantly enriched with both analysis methods, we considered this as sufficient evidence to focus primarily on astrocytes going forward.

Canonical markers GFAP (*p* = 0.00508) and SLC1A3 (*p* = 0.0254) - a glial high affinity glutamate transporter - were among the markers significantly downregulated in depressed OBs in the proteomic analysis while astrocyte-specific markers ALDH1L1, ALDOC, NFIA and GJA1 showed a downregulation trend (Figure 2A). To confirm these observations in situ, at the transcriptional level, with a highly sensitive method, we carried out multiplexed fluorescence in situ hybridization (FISH; RNAScope®) to assess canonical astrocytic marker transcripts. We used the pan-astrocytic marker ALDH1L1 and counted positive cells as cells that expressed both the target transcript and ALDH1L1. These experiments confirmed that ALDOC, SLC1A3 (EAAT1), GJA1 (Connexin 43) and NFIA are primarily expressed in human OB astrocytes (Supplementary Figure 2). Furthermore, the number of ALDH1L1-positive cells was not significantly different between groups (*p* = 0.1406). However, the number of astrocytes expressing ALDOC (*p* = 0.0272), GJA1 (*p* = 0.0021), NFIA (*p* = 0.0003) and SLC1A3 (*p* = 0.0030) were significantly reduced in the OB of depressed individuals compared to controls (Figure2B-F). As ALDOC is a glycolytic enzyme, its downregulation in depression can be related to the reduction of astrocytic glutamate transporters such as SLC1A3, which was also observed in the OB of cases. According to the astrocyte-neuron lactate-shuttle hypothesis, astrocytes upregulate glycolysis to produce lactate for neurons when glutamate is taken up from the synapse (O’Leary and Mechawar, 2021; Pellerin and Magistretti, 1994; Zhao et al., 2016). Given the results obtained with FISH showing no significant group difference in cells expressing ALDH1L1, we examined this further using IF with an antibody against ALDH1L1. As illustrated in Supplementary figure 3, results were consistent, revealing no significant group difference in the density of ALDH1L1-immunoreactive cells in the dorsal (p = 0.556) nor ventral (p = 0.905) OB sections. Similarly, pooling data for both OB regions led to similar densities between cases and controls (p = 0.905).

**Figure 2.**
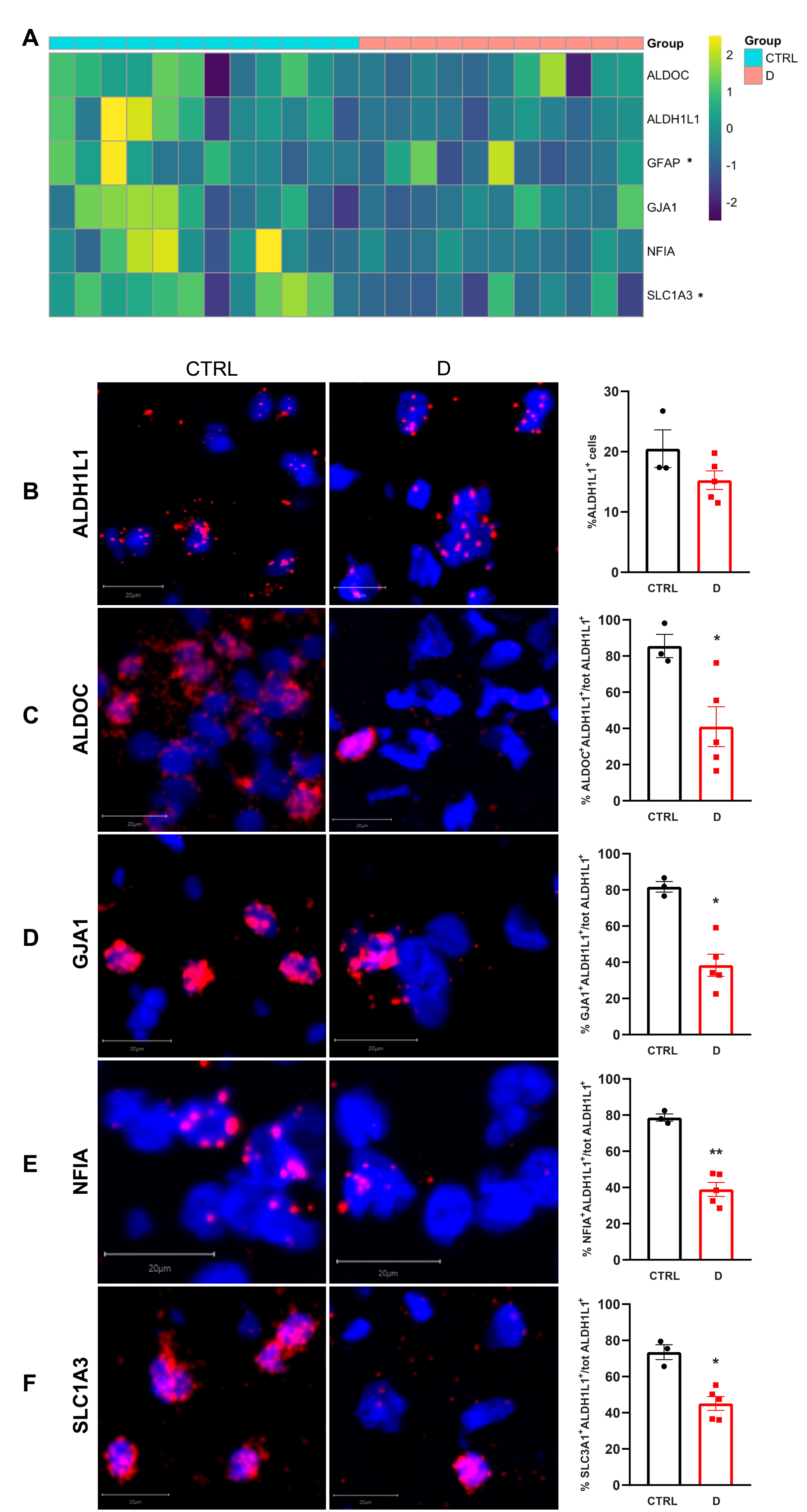
Translational deregulation of canonical astrocytic markers in OBs of depressed individuals. (**A**) Heat map illustrating differentially abundant canonical astrocytic markers in OBs from CTRL individuals as opposed to those from depressed subjects. GFAP and SLC1A3 proteins were significantly downregulated in MDD OBs (respectively, *p* = 0.0013 and *p* = 0.017), while ALDOC, NFIA, ALDH1L1, and GJA1 showed a downregulation trend. (**B**) Left: Representative microphotographs of FISH experiments on CTRL and MDD OBs using ALDH1L1 probe. Right: percentage of ALDH1L1^+^ cells, *p* = 0.1406. **C-F**) Left: Representative microphotographs of FISH experiments on control (CTRL) and depressed patients (D) OBs using ALDH1L1 probe along with (**C**) ALDOC, (**D**) GJA1, (**E**) NFIA, or (**F**) SLC1A3 probes. Right: percentage of double positive astrocytes over the total number of astrocytes (ALDH1L1^+^ cells). (**C**) *p* = 0.0272; (**D**) *p* = 0.0021; (**E**) *p* = 0.0003; (**F**) *p* = 0.0030. (**B**-**F**) CTRL: n = 3; MDD: n = 5; scale bars: 20µm.

We then accessed a single nucleus RNA sequencing (snRNA-seq) dataset recently generated by our group with human dorsolateral PFC (dlPFC) from depressed suicides and control adult males (Maitra et al., 2023) to compare the differentially expressed genes in its astrocyte clusters to our proteome results. Despite differences in brain regions, we found overlapping genes between the astrocyte 1 and 2 dlPFC cluster from the Maitra et al. study (in a re-analysis of the male data from Nagy et al., 2020) and the proteomics data presented here, with all 8 overlapping genes displaying a downregulation compared to controls, according to log fold change (Supplemental Table 7) which indicates a strong concordance between RNA and protein changes (Maitra et al., 2023; Nagy et al., 2020). Some of these proteins, such as AQP1, MT1X and S1PR1, are highly expressed by astrocytes, and elicit crucial roles in homeostatic and immune-related functions. Namely, S1PR1 is implicated in proliferation and trafficking of immune cells (Rothhammer et al., 2017). By generating high levels of antioxidants including metallothionein-I (MT1X), astrocytes play a key role in offsetting oxidative stress associated with inflammation (Waller et al., 2018).

We then verified that these depression-associated changes in OB astrocytic proteins were not driven by confounders such as child abuse history and antidepressant treatment. As illustrated in Supplementary Figure 4, no significant relationship was found between protein quantities and either of these potential confounders using the Mann-Whitney U-Test (child abuse (CA): ALDOC, *p* = 0.456; ALDH1L1, *p* = 0.611; GFAP, *p* = 0.2505; GJA1, *p* = 0.845; NFIA, *p* = 0.785; SLC1A3, *p* = 0.667. Antidepressant treatment: ALDOC, *p* = 0.413; ALDH1L1, *p* = 0.928; GFAP, *p* = 0.133; GJA1, *p* = 0.413; NFIA, *p* = 0.880; SLC1A3, *p* = 0.316).

### Dysregulation of phosphoproteins in the OB of depressed individuals

The phosphoproteome is more dynamic than the proteome, thus we expected some discordance with the proteome results. We found 57 differentially abundant phosphopeptides, corresponding to 50 individual proteins, all of which met the 2-sigma fold change cut-off (Figure 3A). Gene ontology indicated that the corresponding proteins of the differentially abundant phosphopeptides are involved in biological processes such as synaptic vesicle cycle (e.g., AMPH, SYN1, SYN2, PPFIA2), phosphorus metabolic processes (e.g., PNPO, ATP2B4, SLC4A1), and RNA splicing (e.g., TRA2A, SRSF1, SRRM1, THRAP3) (Figure 3B). We assessed overlap between this list of 57 phosphopeptides and DisGeNET differentially expressed genes for depression and found 10 overlapping candidates (Figure 3C). We performed the same cell-type specific hypergeometric tests as in the proteomic analysis and that both neurons (*p* = 0.0125), and astrocytes (*p* = 0.0281) were significantly enriched, with neurons having the highest enrichment score (endothelial, *p* = 0.665; microglia, *p* = 1.000; OL-lineage, *p* = 0.733; Figure 3D).

**Figure 3.**
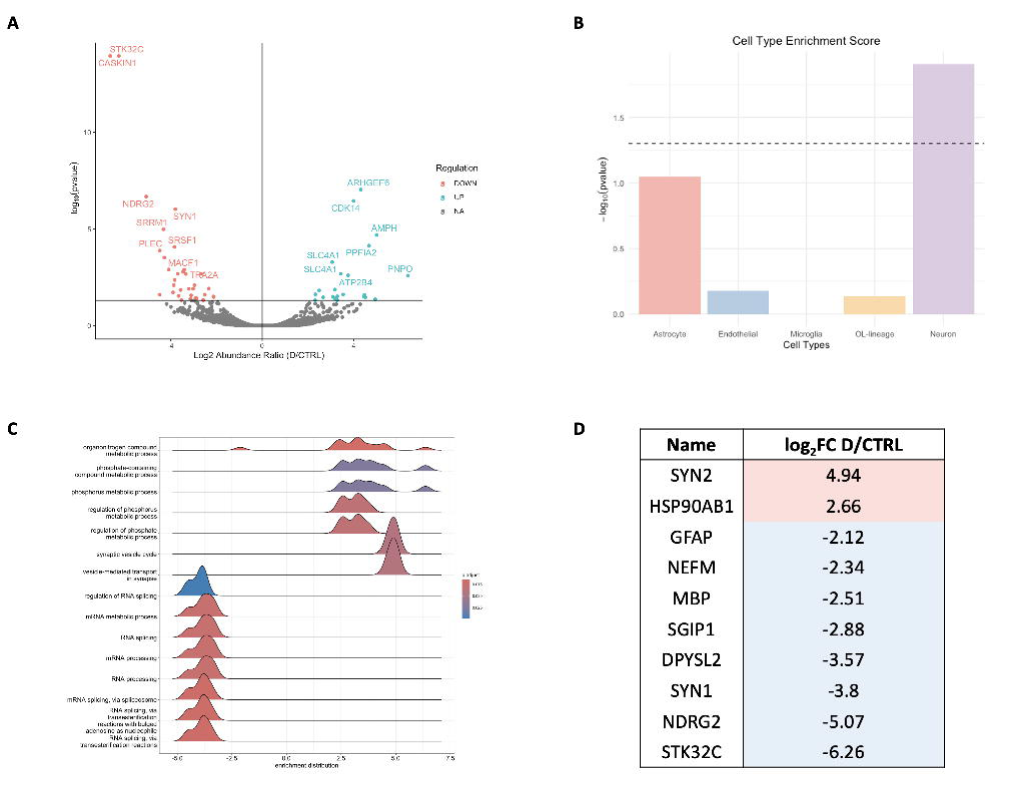
Human OB differential phosphoproteomics in depressed individuals. (**A**) Volcano plot with log base 2 abundance ratio between cases and controls on the x-axis and the -log_10_(p- value) on the y-axis. Each dot represents a protein that showed significant phosphopeptide differential regulation between cases and controls, and the points are colored by direction of the regulation of the abundance of the phosphopeptide modification with downregulation in pink and upregulation in turquoise. The log base 2 abundance ratio cutoff is |3| for visualization purposes and the p-value is cutoff is -log_10_(0.05). D = Depression, CTRL = Control. (**B**) Ridge plot with the enrichment of the gene ontology (GO) terms on the x-axis, and each GO biological process listed on the y-axis. This plot shows the over enriched GO terms in the list of differentially abundant proteins between cases and controls. The ridges are colored by adjusted p-value of the enrichment tests as demonstrated by the color bar. The positive enrichment distribution values indicate enrichment of the GO term in cases compared to controls, and the negative enrichment distribution values indicate enrichment in controls compared to cases (**C**) Table of overlap of proteins in list of proteins with differentially abundant phosphopeptide and genes found in the DisGeNET database. Proteins are listed in descending order by log base 2-fold change between cases and controls. Abundance of the differentially abundant phosphopeptide modification for the protein between cases and controls is represented by log base 2-fold change, phosphopeptide with positive fold change values are colored in pink and negative fold change values are colored in blue. In cases where there were multiple phosphopeptide regulations for a given protein, the average fold change across all modifications was used. (**D**) Bar plot showing cell type enrichment scores of differentially abundant proteins. The x-axis shows the broad cell types of the brain, and the y-axis shows the -log_10_(p-value), where the p-value refers to the enrichment p-value from hypergeometric tests. The dotted line refers to the significance cutoff which is - log_10_(0.05). Only the neurons and astrocytes surpass this significance threshold.

We also examined the relationship between the proteome and phosphoproteome differential abundance. As listed in Supplemental Table 8, there are eight proteins overlapping between the two analyses, including GFAP and AQP1, which are fairly specific to astrocytes in the adult human brain. SLC1A4 had two differentially abundant phosphopeptides, both upregulated, as was observed for the protein. In fact, the regulation direction was found to be consistent for all proteins. For instance, both GFAP protein and a GFAP phosphopeptide were downregulated in cases compared to controls and was identified in the DisGeNET dataset as well.

### Dysregulation of oligodendrocyte proteins in OB of socially defeated mice

To examine whether human OB findings could be observed in a mouse model of depression, we performed the same proteomics analyses with socially defeated mice and controls. Proteins were extracted from 3 control and 3 socially-defeated OB pairs (each sample was derived from 2 OBs, yielding 6 subjects per group in total). As presented in the Methods section, only susceptible mice consistently showing susceptibility throughout the behavioral tests (Open Field, Social Interaction and Elevated Plus Maze) following chronic stress were chosen for proteomic analyses (Figure 4A-C). The analysis revealed 261 significantly DAPs, 193 of which also met the fold change cut-off (Figure 4D). Gene ontology analyses suggested that these proteins are involved in functions such as fatty acid biosynthetic process (e.g., ptgds, anxa1, acsm3) and neurogenesis (e.g. snapin, ndel1, cit) (Figure 4E). We found 35 DAPs overlapping with the DisGeNET database, indicating overlap with genes previously implicated in MDD (Figure 4F). Strikingly, we found a different pattern of cell type enrichment in the DAPs, as the only cell type reaching our significance cut-off was OL-lineage cells (*p* = 0.0453), with endothelial cells showing a trend (endothelial *p* = 0.0679, astrocyte *p* = 0.733, microglia *p* = 0.627, neuronal *p* = 0.375) (Figure 4G). For instance, plp1 (*p* = 0.0106), anln (*p* = 0.0299), apc (*p* = 0.000765), ptgds (*p* = 0.00000309) and tf (*p* = 0.000000189) were among the strongly dysregulated OL markers in socially defeated mice. These genes have all been implicated in the process of myelination, with plp1 being the most abundant protein in myelin of the central nervous system (Jahn et al., 2009). We then conducted a species comparison and found 13 overlapping DAPs including cat (involved in oxidative stress) and apoa1 (involved in lipid metabolism). According to log fold change, 11 of these 13 proteins were regulated in the same direction in both species (Supplemental Table 9).

**Figure 4.**
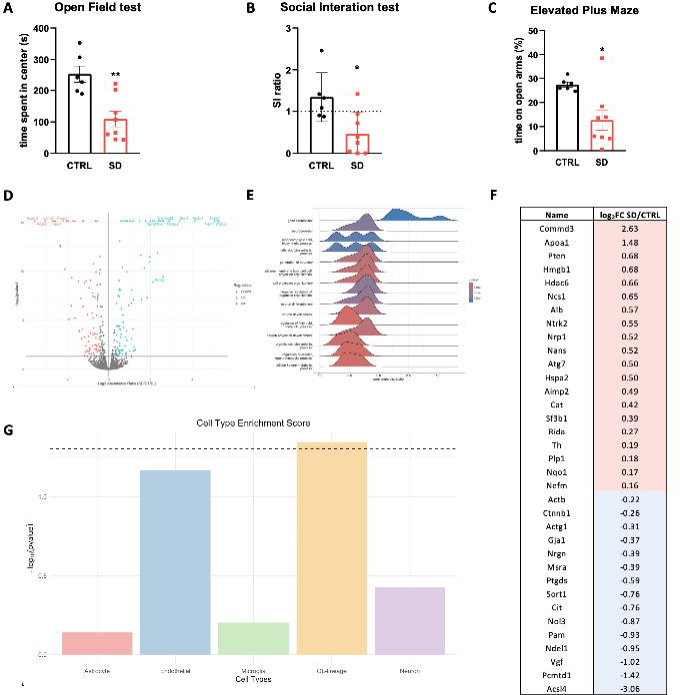
OB proteomics of social defeat stress mice do not recapitulate human OB proteomics of depressed individuals. Plots **A**-**C** demonstrate the behavioral tests conducted on control and socially defeated mice, CTRL, n = 6; SD, n = 8. (**A**) *p* = 0.002; (**B**) *p* = 0.012; (**C**) *p* = 0.012. (**D**) Volcano plot with log base 2 abundance ratio between cases and controls on the x- axis and the -log_10_(p-value) on the y-axis. Each dot represents a protein, and they are colored by direction of regulation with downregulation in pink and upregulation in turquoise. The log base 2 abundance ratio cutoff is |2.5| for visualization purposes and the p-value is cutoff is -log_10_(0.05). (**E**) Ridge plot with the enrichment of the gene ontology (GO) terms on the x-axis, and each GO biological process listed on the y-axis. This plot shows the over enriched GO terms in the list of differentially abundant proteins between cases and controls. The ridges are colored by adjusted p-value of the enrichment tests as demonstrated by the color bar. The positive enrichment distribution values indicate enrichment of the GO term in SD compared to controls, and the negative enrichment distribution values indicate enrichment in controls compared to SD. (**F**) Table of overlap of proteins in list of differentially abundant proteins and genes found in the DisGeNET database. Proteins are listed in descending order by log base 2-fold change between cases and controls. Proteins with positive fold change values are colored in pink and negative fold change values are colored in blue. (**G**) Bar plot showing cell type enrichment scores of differentially abundant proteins. The x-axis shows the broad cell types of the brain, and the y- axis shows the -log_10_(p-value), where the p-value refers to the enrichment p-value from hypergeometric tests. The dotted line refers to the significance cutoff which is -log_10_(0.05). Only the OL-lineage cells surpass this significance threshold.

## DISCUSSION

This is the first study to compare adult OB proteomic and phosphoproteomic profiles between depressed individuals and healthy controls. We observed that depressed OBs presented DAPs that are enriched for astrocyte markers, which led us to explore whether these findings were reflective of changes in the astrocyte population. While the percentage of ALDH1L1^+^ astrocytes remained similar between cases and controls, the percentages of astrocytes expressing the canonical markers ALDOC, GJA1, NFIA, and SLC1A3 were significantly reduced in the OB of cases. In addition to these cell type-enriched changes, we also found that there was a substantial overlap between depression-associated dysregulated OB proteins and genes previously implicated in depression across other brain regions. Moreover, parallel analyses conducted with a mouse model of depression revealed that, in contrast to humans, differentially abundant proteins in the OB of socially stressed animals were enriched for oligodendrocyte-lineage markers.

Astrocytic abnormalities have been consistently implicated in depression and suicide (Torres-Platas et al., 2016), with reported downregulations of astrocyte-specific genes, decreased immunoreactivity to astrocyte-canonical markers, and reduced blood vessel coverage by astrocytic endfeet in postmortem brain samples from depressed patients (Rajkowska et al., 2013). Surprisingly, and despite neuroimaging and psychometric studies implicating the OB in depression, this major limbic region had never been examined at the cellular/molecular level in depressed patients. Our results provide the first evidence that in depression, astrocytic abnormalities are present in the OB, in agreement with our previous reports of astrocytic abnormalities in other cortico-limbic brain regions of depressed suicides (O’Leary et al., 2021; O’Leary and Mechawar, 2021; Torres-Platas et al., 2016; Xiong et al., 2019), as well as with independent animal studies having implicated astrocytes in depression-like behavior (Abbink et al., 2020; Codeluppi et al., 2021; Du Preez et al., 2021; Rurak et al., 2021; Simard et al., 2018).

Depression-associated changes in astrocytic numbers and functions could result from a variety factors, such as chronic glucocorticoid exposure or altered neurotrophic or neurotransmitter signaling (Rajkowska and Miguel-Hidalgo, 2007). The current study indirectly implicates the role of astrocytic transcription factor NFIA, which has been reported to regulate astrocyte morphology and function in the rodent hippocampus (Huang et al., 2020). In addition, these authors found that NFIA-deficient astrocytes or simply hypo-functional NFIA can impair neuron-astrocyte communication *in vitro* (Huang et al., 2020). Interestingly, within 5 days, transient NFIA expression *in vitro* activates glial potential in human pluripotent stem cells, transforming them into astrocytes upon exposure to glial-promoting factors. These NFIA- induced astrocytes not only foster synaptogenesis but also exhibit neuroprotective properties (Tchieu et al., 2019). We can speculate that the reduction in NFIA measured here in the OB of depressed individuals impairs the morpho-functional properties of astrocytes, having in turn an impact on OB volume and olfactory sensitivity, which have both been documented in depressed patients (Negoias et al., 2010; Pause et al., 2001). A smaller OB volume has been associated with higher rates of depressive episodes and with decreased signaling with other limbic areas (Athanassi et al., 2021; Croy and Hummel, 2017). Altered olfaction in depression has been also hypothesized to be linked to a decreased receptor turnover rate in the olfactory epithelium (Athanassi et al., 2021; Croy and Hummel, 2017). Our finding of astrocytic disruptions in the OB of depressed individuals might be associated with this phenomenon and contribute directly to changes in perception, since mature OB astrocytes, at least in mice, are required for normal physiological sensory processing (Ung et al., 2021).

To better understand the effects of protein regulation in depression, our study also compared differentially phosphorylated proteins in OB samples from depressed patients compared to healthy controls. Changes in phosphorylation almost always reflects a change in protein activity, although many such modifications remain uncharacterized (Ardito et al., 2017). In agreement with our DAP analysis, which highlighted an enrichment of astrocyte proteins in the OB of depressed individuals, the phosphoproteomic analysis revealed a corresponding enrichment in differentially phosphorylated astrocytic proteins. Interestingly, it also revealed enrichment in differentially phosphorylated neuronal proteins. It is well established that astrocytes are an essential component of synapses, regulating both synaptic and circuit plasticity and function (Perez-Catalan et al., 2021). In consequence, the astrocytic dysregulation suggested by our study may impact neuronal functions at various levels, including through post-translational modifications of synaptic proteins. Furthermore, given the converging evidence implicating alterations in synaptic density, synapse microenvironment, and synaptic vesicle release (Duman et al., 2016; Holmes et al., 2019; Ren and Guo, 2021) in depression, it is not surprising that some of our most prominent differentially abundant phosphoproteins are localized at the synapse. This can be illustrated by synapsin (SYN) 1 and 2, which were found to be significantly different between cases and controls. Synapsins are ubiquitous phosphoproteins on synaptic vesicles, where they regulate the trafficking of vesicles in nerve terminals, a critical parameter in short-term synaptic plasticity (Sun et al., 2006). A previous study of depression in postmortem dlPFC from males and females pointed to synaptic function as the main biological process associated with differentially abundant phosphoproteins (Martins-de-Souza et al., 2012). Interestingly, 10 out of the 50 phosphoproteins we found to be differentially abundant were similarly differentially abundant in the latter study, despite the regional difference. In our results, SYN 1 was one of the top differentially abundant phosphoproteins with three significantly downregulated phosphopeptides, indicating hypophosphorylation of SYN 1 in depressed individuals compared to controls. This finding was mirrored by significantly downregulated SYN 1 phosphopetides in Martins-de-Souza et al. (2012) (Martins-de-Souza et al., 2012). The sizable overlap between these two studies further strengthens the evidence for post-translational modifications occurring at the synapse in depression.

We then conducted the same OB proteomic analysis in a mouse model of depression. These experiments in the social defeat mouse model indicated that the enriched astrocyte signature observed in humans with depression is not shared by chronically stressed mice, possibly highlighting a limitation of this model which recapitulates many of the molecular signatures of depression in regions such as the medial prefrontal cortex (mPFC) and nucleus accumbens (Scarpa et al., 2020).

The social defeat model of human depression comes with some limitations. This model involves daily defeat and thus fights, which involve stress, wounds, and local inflammatory events. The model also differs in many ways from the post-mortem human brains from depressed patients. Namely, depression-like states in mice appear on vastly different timescales than those of disease processes implicated in depressive symptoms in humans. Moreover, there exist differences in pathological and adaptive mechanisms when comparing this animal model and depressed humans. Finally, social defeat is performed almost exclusively in adult animals, which cannot inform on the contribution of early-life events to the depression in adulthood.

Our observation adds to the extensive breadth of evidence supporting OL lineage abnormalities that have been reported in post-mortem and animal model studies of depression (Liu et al., 2012; Makinodan et al., 2012; Nagy et al., 2020; Rahimian et al., 2022). In a study of chronic social defeat stress, decreases in myelin proteins were observed in the nucleus accumbens and mPFC, with the mPFC also displaying thinner myelin and decreased internode length (Bonnefil et al., 2019). In a subsequent study, this model led long lasting reductions of OL precursor cells (OPC), abnormal differentiation rates of OPC to OLs, and extensive hypomyelination (Kokkosis et al., 2022). In fact, these findings are not limited to social defeat stress, as myelin-related molecular and ultrastructural changes have been reported in several models of stress including social isolation and chronic variable stress (Bonnefil et al., 2019; Liu et al., 2012; Rahimian et al., 2022). While our OL enrichment findings may point to differences in myelin content or regulation in socially defeated mice, further experiments are needed to better understand the biological significance of these results. It is also a possibility that discordance in results might reflect differences in OB cellular composition between species. In summary, while there was overlap between human and mouse DAPs, as well as an overlap with previous findings from DisGeNET, the cell type enrichment and gene ontology analyses indicated considerable divergence between results in humans and mice, indicating possible mechanistic differences between the murine social defeat model and human depression, at least in what concerns the OB. It is plausible to assume that this divergence is at least partially related to the disproportionately greater OB/brain volume in mice compared to humans.

The current study contributes to a better understanding of the cellular and molecular factors associated with depression, which is required for the design of impactful translational studies, including research on diagnostic or prognostic biomarkers as well as on emerging therapeutics. A growing body of evidence suggests that astrocyte-derived extracellular vesicles can be obtained non-invasively from the serum of clinically depressed patients and inform on the functional state of astrocytes *in vivo* (Xie et al., 2023). It remains to be determined if peripheral levels of these proteins are correlated with astrocytic deficits in one or multiple regions of the CNS and with one or more astrocytic subtypes.

The main limitations of our study were its sample size and the fact that all the OBs were from male donors. It is challenging to collect postmortem OB samples, and particularly samples that are of sufficient quality for proteomics, and this becomes even more difficult by the very low number of OBs collected from female middle-aged depressed individuals over the years. As a result, we were unable to include female samples in a meaningful way. Given the distinct male vs female molecular signatures of depression that have been reported in other brain regions, especially in astrocytes and microglia (Maitra et al., 2023) and the sexual dimorphism in OB cellular composition in humans (Oliveira-Pinto et al., 2014), future investigations will need to circumvent these challenges to explore whether the OB is similarly affected in females affected by depression. Lastly, since all but one of the cases included in this study died by suicide, we cannot exclude that the proteomic signature reported in this study is more representative of male depressed suicides than of all depressed individuals (Peng et al., 2023).

Overall, the findings of our study further implicate astrocytes in depression. The consistency of cortico-limbic astrocyte findings in depression research lends support to the notion that this cell type might play a key role in the pathophysiology of this heterogeneous illness. Furthermore, our study raises the possibility that the chronic social defeat stress model of depressive-like behavior might not sufficiently recapitulate the molecular neurobiology of human depression and suicide, at least in the OB. Understanding the strengths and weaknesses of animal models of depression will help to better position pre-clinical research questions such that their translatability to human depression may be enhanced.

## Data availability

The mass spectrometry proteomics data have been deposited to the ProteomeXchange Consortium via the PRIDE partner repository with the dataset identifier PXD045011

**Project Name:** Proteomic evidence of depression-associated astrocytic dysfunction in the human male olfactory bulb

**Project accession:** PXD045011

The processed spreadsheets for human proteomics and phosphoproteomics, as well as the mouse proteomics have been provided as Supplementary Tables 10 (human proteomics). Supplementary Table 11 (human phosphoproteomics), and Supplementary Table 12 (mouse proteomics). The cell type marker list for the enrichment analysis has been as Supplementary Table 6 as well.

## AUTHOR CONTRIBUTIONS

NM conceptualized the study. RR prepared samples for proteomic experiments and performed all FISH validations. RR, KP and NM wrote the manuscript and prepared the figures. KP performed the bioinformatic analyses for the proteomic and phosphoproteomic data. RM and MAD conducted histological work. GF and LP performed animal behavioral tests. GF extracted mouse OBs and analyzed the behavioral tasks. CH contributed to the FISH analysis. VR, RPZ and CB performed proteomic and phosphoproteomic experiments. NM, BG, CN and GT supervised the project. All authors read, contributed to, and agreed with the final version of the manuscript.

### ACKNOWLEDGMENTS

This study was funded by an ERA NET Neuron grant and a CIHR Project grant to N.M. R.R. and G.F. received fellowships and K.P. a scholarship from the FRQ-S. K.P. currently holds a CIHR doctoral scholarship and R.M. a scholarship from the Botswana government. The Douglas-Bell Canada Brain Bank is supported in part by platform support grants from the Réseau Québécois sur le Suicide, les Troubles de l’Humeur et les Troubles Associés (FRQ-S), Healthy Brains for Healthy Lives (CFREF), and Brain Canada. This study used the services of the Molecular and Cellular Microscopy Platform (MCMP) at the Douglas Research Centre.

This work was also supported by funding to CHB from Genome Canada’s Genomics Technology Platform (264PRO). CHB is also grateful for support from the Segal McGill Chair in Molecular Oncology at McGill University (Montreal, Quebec, Canada) for support from the Terry Fox Research Institute, the Warren Y. Soper Charitable Trust, and the Alvin Segal Family Foundation to the Jewish General Hospital (Montreal, Quebec, Canada).

This work was done under the auspices of a Memorandum of Understanding between McGill and the U.S. National Cancer Institute’s International Cancer Proteogenome Consortium (ICPC). ICPC encourages international cooperation among institutions and nations in proteogenomic cancer research in which proteogenomic datasets are made available to the public. This work was also done in collaboration with the U.S. National Cancer Institute’s Clinical Proteomic Tumor Analysis Consortium (CPTAC).

Lastly, the authors wish to thank the donors and their families for their invaluable gift.

## ADDITIONAL INFORMATION

Supplementary information

## REFERENCES

1. Abbink, M.R., Kotah, J.M., Hoeijmakers, L., Mak, A., Yvon-Durocher, G., van der Gaag, B., Lucassen, P.J., Korosi, A., 2020. Characterization of astrocytes throughout life in wildtype and APP/PS1 mice after early-life stress exposure. J. Neuroinflammation 17, 91.

2. Ardito, F., Giuliani, M., Perrone, D., Troiano, G., Lo Muzio, L., 2017. The crucial role of protein phosphorylation in cell signaling and its use as targeted therapy (Review). Int. J. Mol. Med. 40, 271–280.

3. Athanassi, A., Dorado Doncel, R., Bath, K.G., Mandairon, N., 2021. Relationship between depression and olfactory sensory function: a review. Chem. Senses 46.

4. Beck, F., Geiger, J., Gambaryan, S., Solari, F.A., Dell’Aica, M., Loroch, S., Mattheij, N.J., Mindukshev, I., Potz, O., Jurk, K., Burkhart, J.M., Fufezan, C., Heemskerk, J.W., Walter, U., Zahedi, R.P., Sickmann, A., 2017. Temporal quantitative phosphoproteomics of ADP stimulation reveals novel central nodes in platelet activation and inhibition. Blood 129, e1–e12.

5. Bonnefil, V., Dietz, K., Amatruda, M., Wentling, M., Aubry, A.V., Dupree, J.L., Temple, G., Park, H.J., Burghardt, N.S., Casaccia, P., Liu, J., 2019. Region-specific myelin differences define behavioral consequences of chronic social defeat stress in mice. Elife 8.

6. Burkhart, J.M., Schumbrutzki, C., Wortelkamp, S., Sickmann, A., Zahedi, R.P., 2012. Systematic and quantitative comparison of digest efficiency and specificity reveals the impact of trypsin quality on MS-based proteomics. J. Proteomics 75, 1454–1462.

7. Codeluppi, S.A., Chatterjee, D., Prevot, T.D., Bansal, Y., Misquitta, K.A., Sibille, E., Banasr, M., 2021. Chronic Stress Alters Astrocyte Morphology in Mouse Prefrontal Cortex. Int. J. Neuropsychopharmacol. 24, 842–853.

8. Croy, I., Hummel, T., 2017. Olfaction as a marker for depression. J. Neurol. 264, 631–638.

9. Croy, I., Negoias, S., Novakova, L., Landis, B.N., Hummel, T., 2012. Learning about the functions of the olfactory system from people without a sense of smell. PLoS One 7, e33365.

10. Du Preez, A., Onorato, D., Eiben, I., Musaelyan, K., Egeland, M., Zunszain, P.A., Fernandes, C., Thuret, S., Pariante, C.M., 2021. Chronic stress followed by social isolation promotes depressive-like behaviour, alters microglial and astrocyte biology and reduces hippocampal neurogenesis in male mice. Brain. Behav. Immun. 91, 24–47.

11. Dumais, A., Lesage, A.D., Lalovic, A., Seguin, M., Tousignant, M., Chawky, N., Turecki, G., 2005. Is violent method of suicide a behavioral marker of lifetime aggression? Am. J. Psychiatry 162, 1375–1378.

12. Duman, R.S., Aghajanian, G.K., Sanacora, G., Krystal, J.H., 2016. Synaptic plasticity and depression: new insights from stress and rapid-acting antidepressants. Nat. Med. 22, 238–249.

13. Golden, S.A., Covington, H.E., 3rd, Berton, O., Russo, S.J., 2011. A standardized protocol for repeated social defeat stress in mice. Nat. Protoc. 6, 1183-1191.

14. Gonczarowska-Jorge, H., Loroch, S., Dell’Aica, M., Sickmann, A., Roos, A., Zahedi, R.P., 2017. Quantifying missing (phospho)proteome regions with the broad-specificity protease subtilisin. Anal. Chem. 89, 13137–13145.

15. Holmes, S.E., Scheinost, D., Finnema, S.J., Naganawa, M., Davis, M.T., DellaGioia, N., Nabulsi, N., Matuskey, D., Angarita, G.A., Pietrzak, R.H., Duman, R.S., Sanacora, G., Krystal, J.H., Carson, R.E., Esterlis, I., 2019. Lower synaptic density is associated with depression severity and network alterations. Nat Commun 10, 1529.

16. Huang, A.Y., Woo, J., Sardar, D., Lozzi, B., Bosquez Huerta, N.A., Lin, C.J., Felice, D., Jain, A., Paulucci-Holthauzen, A., Deneen, B., 2020. Region-Specific Transcriptional Control of Astrocyte Function Oversees Local Circuit Activities. Neuron 106, 992–1008 e1009.

17. Jahn, O., Tenzer, S., Werner, H.B., 2009. Myelin proteomics: molecular anatomy of an insulating sheath. Mol. Neurobiol. 40, 55–72.

18. Kelly, J.P., Wrynn, A.S., Leonard, B.E., 1997. The olfactory bulbectomized rat as a model of depression: an update. Pharmacol. Ther. 74, 299–316.

19. Kohli, P., Soler, Z.M., Nguyen, S.A., Muus, J.S., Schlosser, R.J., 2016. The Association Between Olfaction and Depression: A Systematic Review. Chem. Senses 41, 479–486.

20. Kokkosis, A.G., Madeira, M.M., Mullahy, M.R., Tsirka, S.E., 2022. Chronic stress disrupts the homeostasis and progeny progression of oligodendroglial lineage cells, associating immune oligodendrocytes with prefrontal cortex hypomyelination. Mol. Psychiatry 27, 2833–2848.

21. Korotkevich, G., Sukhov, V., Budin, N., Shpak, B., Artyomov, M.N., Sergushichev, A., 2021. Fast gene set enrichment analysis. bioRxiv, 060012.

22. Liu, J., Dietz, K., DeLoyht, J.M., Pedre, X., Kelkar, D., Kaur, J., Vialou, V., Lobo, M.K., Dietz, D.M., Nestler, E.J., Dupree, J., Casaccia, P., 2012. Impaired adult myelination in the prefrontal cortex of socially isolated mice. Nat. Neurosci. 15, 1621–1623.

23. Lombion-Pouthier, S., Vandel, P., Nezelof, S., Haffen, E., Millot, J.L., 2006. Odor perception in patients with mood disorders. J. Affect. Disord. 90, 187–191.

24. Maitra, M., Mitsuhashi, H., Rahimian, R., Chawla, A., Yang, J., Fiori, L.M., Davoli, M.A., Perlman, K., Aouabed, Z., Mash, D.C., Suderman, M., Mechawar, N., Turecki, G., Nagy, C., 2023. Cell type specific transcriptomic differences in depression show similar patterns between males and females but implicate distinct cell types and genes. Nat Commun 14, 2912.

25. Makinodan, M., Rosen, K.M., Ito, S., Corfas, G., 2012. A critical period for social experience-dependent oligodendrocyte maturation and myelination. Science 337, 1357–1360.

26. Martins-de-Souza, D., Guest, P.C., Vanattou-Saifoudine, N., Rahmoune, H., Bahn, S., 2012. Phosphoproteomic differences in major depressive disorder postmortem brains indicate effects on synaptic function. Eur. Arch. Psychiatry Clin. Neurosci. 262, 657–666.

27. McKenzie, A.T., Wang, M., Hauberg, M.E., Fullard, J.F., Kozlenkov, A., Keenan, A., Hurd, Y.L., Dracheva, S., Casaccia, P., Roussos, P., Zhang, B., 2018. Brain cell type specific gene expression and co-expression network architectures. Sci. Rep. 8, 8868.

28. Milardi, D., Cacciola, A., Calamuneri, A., Ghilardi, M.F., Caminiti, F., Cascio, F., Andronaco, V., Anastasi, G., Mormina, E., Arrigo, A., Bruschetta, D., Quartarone, A., 2017. The Olfactory System Revealed: Non-Invasive Mapping by using Constrained Spherical Deconvolution Tractography in Healthy Humans. Front. Neuroanat. 11, 32.

29. Nagy, C., Maitra, M., Tanti, A., Suderman, M., Theroux, J.F., Davoli, M.A., Perlman, K., Yerko, V., Wang, Y.C., Tripathy, S.J., Pavlidis, P., Mechawar, N., Ragoussis, J., Turecki, G., 2020. Single-nucleus transcriptomics of the prefrontal cortex in major depressive disorder implicates oligodendrocyte precursor cells and excitatory neurons. Nat. Neurosci. 23, 771–781.

30. Navarro, P., Trevisan-Herraz, M., Bonzon-Kulichenko, E., Nunez, E., Martinez-Acedo, P., Perez-Hernandez, D., Jorge, I., Mesa, R., Calvo, E., Carrascal, M., Hernaez, M.L., Garcia, F., Barcena, J.A., Ashman, K., Abian, J., Gil, C., Redondo, J.M., Vazquez, J., 2014. General statistical framework for quantitative proteomics by stable isotope labeling. J. Proteome Res. 13, 1234–1247.

31. Negoias, S., Croy, I., Gerber, J., Puschmann, S., Petrowski, K., Joraschky, P., Hummel, T., 2010. Reduced olfactory bulb volume and olfactory sensitivity in patients with acute major depression. Neuroscience 169, 415-421.

32. O’Leary, L.A., Belliveau, C., Davoli, M.A., Ma, J.C., Tanti, A., Turecki, G., Mechawar, N., 2021. Widespread Decrease of Cerebral Vimentin-Immunoreactive Astrocytes in Depressed Suicides. Front Psychiatry 12, 640963.

33. O’Leary, L.A., Mechawar, N., 2021. Implication of cerebral astrocytes in major depression: A review of fine neuroanatomical evidence in humans. Glia 69, 2077–2099.

34. Oliveira-Pinto, A.V., Santos, R.M., Coutinho, R.A., Oliveira, L.M., Santos, G.B., Alho, A.T., Leite, R.E., Farfel, J.M., Suemoto, C.K., Grinberg, L.T., Pasqualucci, C.A., Jacob-Filho, W., Lent, R., 2014. Sexual dimorphism in the human olfactory bulb: females have more neurons and glial cells than males. PLoS One 9, e111733.

35. Pause, B.M., Miranda, A., Goder, R., Aldenhoff, J.B., Ferstl, R., 2001. Reduced olfactory performance in patients with major depression. J. Psychiatr. Res. 35, 271–277.

36. Pellerin, L., Magistretti, P.J., 1994. Glutamate uptake into astrocytes stimulates aerobic glycolysis: a mechanism coupling neuronal activity to glucose utilization. Proc. Natl. Acad. Sci. U. S. A. 91, 10625–10629.

37. Peng, S., Zhou, Y., Xiong, L., Wang, Q., 2023. Identification of novel targets and pathways to distinguish suicide dependent or independent on depression diagnosis. Sci. Rep. 13, 2488.

38. Perez-Catalan, N.A., Doe, C.Q., Ackerman, S.D., 2021. The role of astrocyte-mediated plasticity in neural circuit development and function. Neural Dev 16, 1.

39. Piñero, J., Bravo, À., Queralt-Rosinach, N., Gutiérrez-Sacristán, A., Deu-Pons, J., Centeno, E., García-García, J., Sanz, F., Furlong, L.I., 2017. DisGeNET: a comprehensive platform integrating information on human disease-associated genes and variants. Nucleic Acids Res. 45, D833–d839.

40. Qiu, Y., Wang, J., Lei, J., Roeder, K., 2021. Identification of cell-type-specific marker genes from co-expression patterns in tissue samples. Bioinformatics 37, 3228–3234.

41. Rahimian, R., Perlman, K., Canonne, C., Mechawar, N., 2022. Targeting microglia-oligodendrocyte crosstalk in neurodegenerative and psychiatric disorders. Drug Discov Today 27, 2562–2573.

42. Rajkowska, G., Hughes, J., Stockmeier, C.A., Javier Miguel-Hidalgo, J., Maciag, D., 2013. Coverage of blood vessels by astrocytic endfeet is reduced in major depressive disorder. Biol. Psychiatry 73, 613–621.

43. Rajkowska, G., Miguel-Hidalgo, J.J., 2007. Gliogenesis and glial pathology in depression. CNS Neurol. Disord. Drug Targets 6, 219–233.

44. Reemst, K., Kracht, L., Kotah, J.M., Rahimian, R., van Irsen, A.A.S., Congrains Sotomayor, G., Verboon, L.N., Brouwer, N., Simard, S., Turecki, G., Mechawar, N., Kooistra, S.M., Eggen, B.J.L., Korosi, A., 2022. Early-life stress lastingly impacts microglial transcriptome and function under basal and immune-challenged conditions. Transl Psychiatry 12, 507.

45. Ren, F., Guo, R., 2021. Synaptic Microenvironment in Depressive Disorder: Insights from Synaptic Plasticity. Neuropsychiatr. Dis. Treat. 17, 157–165.

46. Rothhammer, V., Kenison, J.E., Tjon, E., Takenaka, M.C., de Lima, K.A., Borucki, D.M., Chao, C.C., Wilz, A., Blain, M., Healy, L., Antel, J., Quintana, F.J., 2017. Sphingosine 1- phosphate receptor modulation suppresses pathogenic astrocyte activation and chronic progressive CNS inflammation. Proc. Natl. Acad. Sci. U. S. A. 114, 2012–2017.

47. Rottstaedt, F., Weidner, K., Strauss, T., Schellong, J., Kitzler, H., Wolff-Stephan, S., Hummel, T., Croy, I., 2018. Size matters - The olfactory bulb as a marker for depression. J. Affect. Disord. 229, 193–198.

48. Rurak, G.M., Woodside, B., Aguilar-Valles, A., Salmaso, N., 2021. Astroglial cells as neuroendocrine targets in forebrain development: Implications for sex differences in psychiatric disease. Front. Neuroendocrinol. 60, 100897.

49. Scarpa, J.R., Fatma, M., Loh, Y.E., Traore, S.R., Stefan, T., Chen, T.H., Nestler, E.J., Labonte, B., 2020. Shared Transcriptional Signatures in Major Depressive Disorder and Mouse Chronic Stress Models. Biol. Psychiatry 88, 159–168.

50. Simard, S., Coppola, G., Rudyk, C.A., Hayley, S., McQuaid, R.J., Salmaso, N., 2018. Profiling changes in cortical astroglial cells following chronic stress. Neuropsychopharmacology 43, 1961–1971.

51. Song, C., Leonard, B.E., 2005. The olfactory bulbectomised rat as a model of depression. Neurosci. Biobehav. Rev. 29, 627–647.

52. Sun, J., Bronk, P., Liu, X., Han, W., Sudhof, T.C., 2006. Synapsins regulate use-dependent synaptic plasticity in the calyx of Held by a Ca2+/calmodulin-dependent pathway. Proc. Natl. Acad. Sci. U. S. A. 103, 2880–2885.

53. Takahashi, T., Nishikawa, Y., Yucel, M., Whittle, S., Lorenzetti, V., Walterfang, M., Sasabayashi, D., Suzuki, M., Pantelis, C., Allen, N.B., 2016. Olfactory sulcus morphology in patients with current and past major depression. Psychiatry Res Neuroimaging 255, 60–65.

54. Tanti, A., Belliveau, C., Nagy, C., Maitra, M., Denux, F., Perlman, K., Chen, F., Mpai, R., Canonne, C., Théberge, S., McFarquhar, A., Davoli, M.A., Belzung, C., Turecki, G., Mechawar, N., 2022. Child abuse associates with increased recruitment of perineuronal nets in the ventromedial prefrontal cortex: a possible implication of oligodendrocyte progenitor cells. Mol. Psychiatry 27, 1552–1561.

55. Tchieu, J., Calder, E.L., Guttikonda, S.R., Gutzwiller, E.M., Aromolaran, K.A., Steinbeck, J.A., Goldstein, P.A., Studer, L., 2019. NFIA is a gliogenic switch enabling rapid derivation of functional human astrocytes from pluripotent stem cells. Nat. Biotechnol. 37, 267–275.

56. Torres-Platas, S.G., Nagy, C., Wakid, M., Turecki, G., Mechawar, N., 2016. Glial fibrillary acidic protein is differentially expressed across cortical and subcortical regions in healthy brains and downregulated in the thalamus and caudate nucleus of depressed suicides. Mol. Psychiatry 21, 509–515.

57. Ung, K., Huang, T.W., Lozzi, B., Woo, J., Hanson, E., Pekarek, B., Tepe, B., Sardar, D., Cheng, Y.T., Liu, G., Deneen, B., Arenkiel, B.R., 2021. Olfactory bulb astrocytes mediate sensory circuit processing through Sox9 in the mouse brain. Nat Commun 12, 5230.

58. Waller, R., Murphy, M., Garwood, C.J., Jennings, L., Heath, P.R., Chambers, A., Matthews, F.E., Brayne, C., Ince, P.G., Wharton, S.B., Simpson, J.E., Cognitive, F., Ageing Neuropathology Study, G., 2018. Metallothionein-I/II expression associates with the astrocyte DNA damage response and not Alzheimer-type pathology in the aging brain. Glia 66, 2316–2323.

59. Xie, X.H., Lai, W.T., Xu, S.X., Di Forti, M., Zhang, J.Y., Chen, M.M., Yao, L.H., Wang, P., Hao, K.K., Rong, H., 2023. Hyper-inflammation of astrocytes in patients of major depressive disorder: Evidence from serum astrocyte-derived extracellular vesicles. Brain. Behav. Immun. 109, 51–62.

60. Xiong, W., Cao, X., Zeng, Y., Qin, X., Zhu, M., Ren, J., Wu, Z., Huang, Q., Zhang, Y., Wang, M., Chen, L., Turecki, G., Mechawar, N., Chen, W., Yi, G., Zhu, X., 2019. Astrocytic epoxyeicosatrienoic acid signaling in the medial prefrontal cortex modulates depressive-like behaviors. J. Neurosci. 39, 4606–4623.

61. Yu, G., Wang, L.G., Han, Y., He, Q.Y., 2012. clusterProfiler: an R package for comparing biological themes among gene clusters. OMICS 16, 284–287.

62. Yu, G., Wang, L.G., Yan, G.R., He, Q.Y., 2015. DOSE: an R/Bioconductor package for disease ontology semantic and enrichment analysis. Bioinformatics 31, 608–609.

63. Zhao, J., Verwer, R.W., van Wamelen, D.J., Qi, X.R., Gao, S.F., Lucassen, P.J., Swaab, D.F., 2016. Prefrontal changes in the glutamate-glutamine cycle and neuronal/glial glutamate transporters in depression with and without suicide. J. Psychiatr. Res. 82, 8–15.

