## Supplementary material for "Proteomic evidence of depression-associated astrocytic dysfunction in the human male olfactory bulb"

**Supplementary Methods**

Chronic social defeat stress

Animal experiments were conducted with the approval of the Douglas Animal Care Committee. Male C57BL/6 mice were subjected to a social defeat paradigm, consisting of a daily 5-minute defeat with a different CD-1 aggressor mouse on each of 10 consecutive days (Golden et al. 2011). A defeat lasted for either 5 minutes or 10 attacks, whichever occurred first. Defeats occurred in large rat cages with two halves separated by a plexiglass divider. After each defeat, the mouse remained in the aggressor's home cage but separated from the aggressor by a perforated translucent Plexiglas divider. This allowed the mouse to maintain visual, olfactory and auditory interactions with the aggressor. In a different room, unexposed to CD-1 aggressors and defeated mice, control mice were housed in pairs separated by a Plexiglas divider and were moved to a different cage each day (Golden et al., 2011).

Screening of aggressor CD-1 mice

Prior to beginning the defeats, aggressor CD-1 mice were screened over 3 consecutive days, and selected according to their response to a novel juvenile male C57BL/6J mouse that was placed in their home cage. They were selected if they had an attack latency under 60 seconds, consistent attacks over 180 seconds and if they attacked on 2 consecutive days (Golden et al. 2011).

Social Interaction Test

Twenty-four hours after the last defeat, the defeated mice were tested in the social interaction (SI) test. This was carried out in an open field box (45 cm × 45 cm × 45 cm) with a Plexiglas wire mesh enclosure (10 cm wide × 6.5 cm deep × 42 cm high) placed in a section of the box designated as the ‘social interaction zone’ (SIZ). In the first phase of this test, the mouse was placed in the box for 150 seconds with an empty wire mesh enclosure. After this, the mouse was placed back in his home cage for 30 seconds. For the second phase, the mouse was placed in the box for another 150 seconds, but this time with a novel CD-1 aggressor present in a different wire mesh enclosure. Control mice underwent the same procedure as defeated mice. The time the mouse spent in the SIZ during each phase was measured, and this was used to calculate the individual’s SI ratio and determine whether they were resilient or susceptible. SI ratios were calculated by dividing the time spent in the SIZ with an aggressive CD-1 mouse present by the time spent in the SIZ with the aggressor present. Susceptibility was defined by a ratio smaller than 1 – meaning they spent less time in the SIZ when the aggressor was present (Golden et al., 2011). The SI tests were recorded using Anymaze software, and time spent in the SIZ was determined using TopScan tracking software.

Open Field Test

24 hours after the SI test, mice were tested in Open Field (OF) chambers. An Omnitech digiscan activity monitor was used to measure locomotor activity. The movements of the mice were measured for 10 minutes in open field chambers (42 cm × 42 cm × 42 cm) with photocells and plexiglass walls and floors. The data were recorded using VersaMax software (Golden et al. 2011).

Elevated Plus Maze

Twenty-four hours following the OF test, mice were subjected to the elevated plus maze (EPM). EPM is made up of 2 opposing open arms (30 × 5cm) and 2 opposing closed arms (30 × 5 × 11cm) branching out from a center zone (5 cm2), and is 50cm high. Mice were placed in the center zone of the EPM, facing an open arm, and were allowed to explore the maze for 10 minutes. The mouse was considered to be inside an open or closed arm when the center of its body was inside it. The amount of time spent in the open and closed arms of the maze was determined using TopScan tracking software (Golden et al. 2011).

**Supplementary Figure 1- Cell type marker enrichment conducted with differentially expressed proteins obtained through analysis including covariates reveals enrichment in astrocyte-specific markers in OBs from depressed individuals compared to controls.**

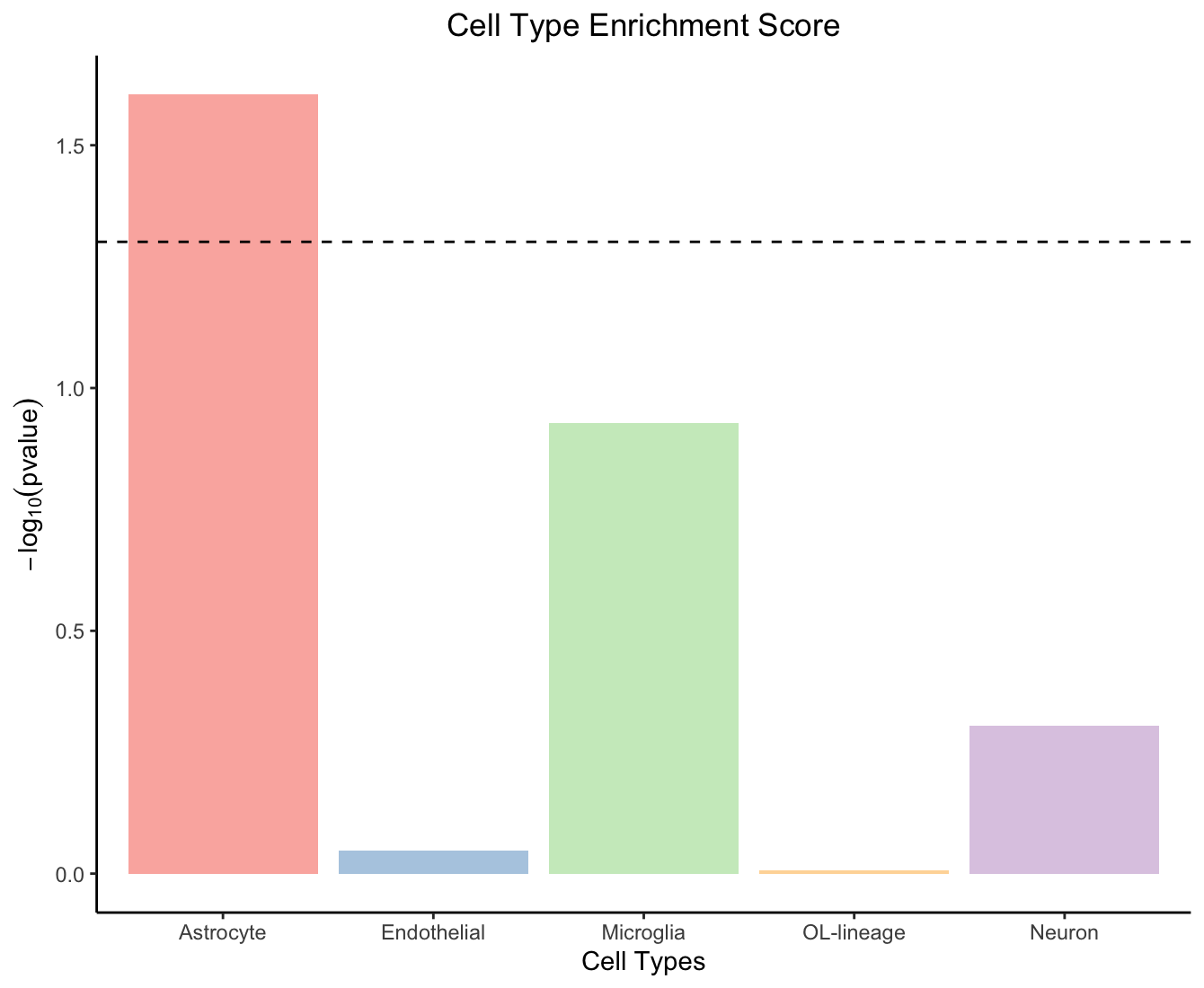

**Supplementary Figure 2. Co-localization of canonical astrocytic marker transcripts with the pan-astrocytic marker ALDH1L1.**

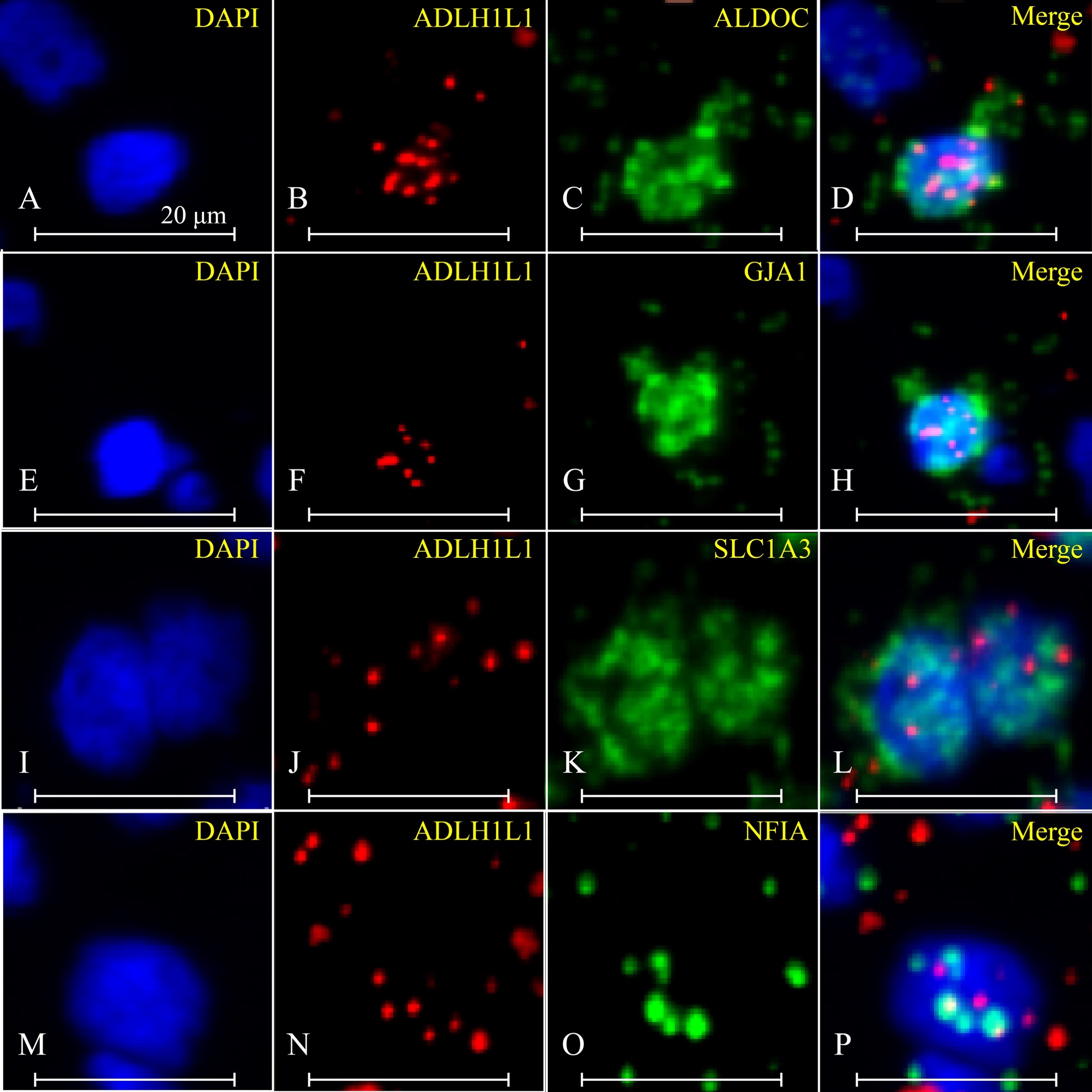

**Supplementary Figure 3. Human olfactory bulb ALDH1L1 immunofluorescence staining.** (A) Representative microphotographs of Immunofluorescence experiments in control (CTRL) and depressed patients (D) using an antibody against ALDH1L1. Left: DAPI (blue). Middle: ALDH1L1 (red). Right: Merge. (B) Density of ALDH1L1^+^ astrocytes pooled between sections from the ventral and dorsal OB; p = 0.905. (C) Density of ALDH1L1^+^ astrocytes in a section sampled from the dorsal OB; p = 0.556. (D) Density of ALDH1L1^+^ astrocytes in a section sampled from the ventral OB; p = 0.905. (A-D) CTRL: n = 4; D: n = 5; scale bars: 20µm

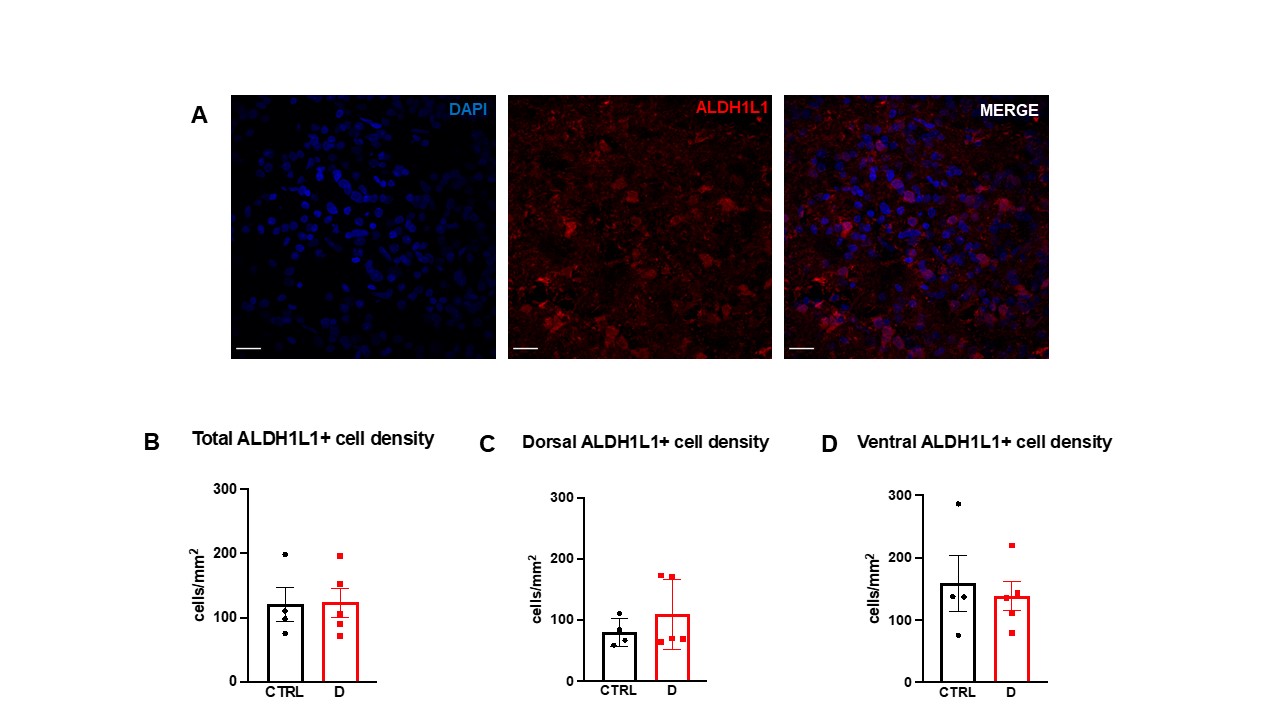

**Supplementary Figure 4. Observed dysregulation of canonical astrocytic markers in the OB of depressed individuals is not driven by a history of child abuse nor by antidepressant treatment.**

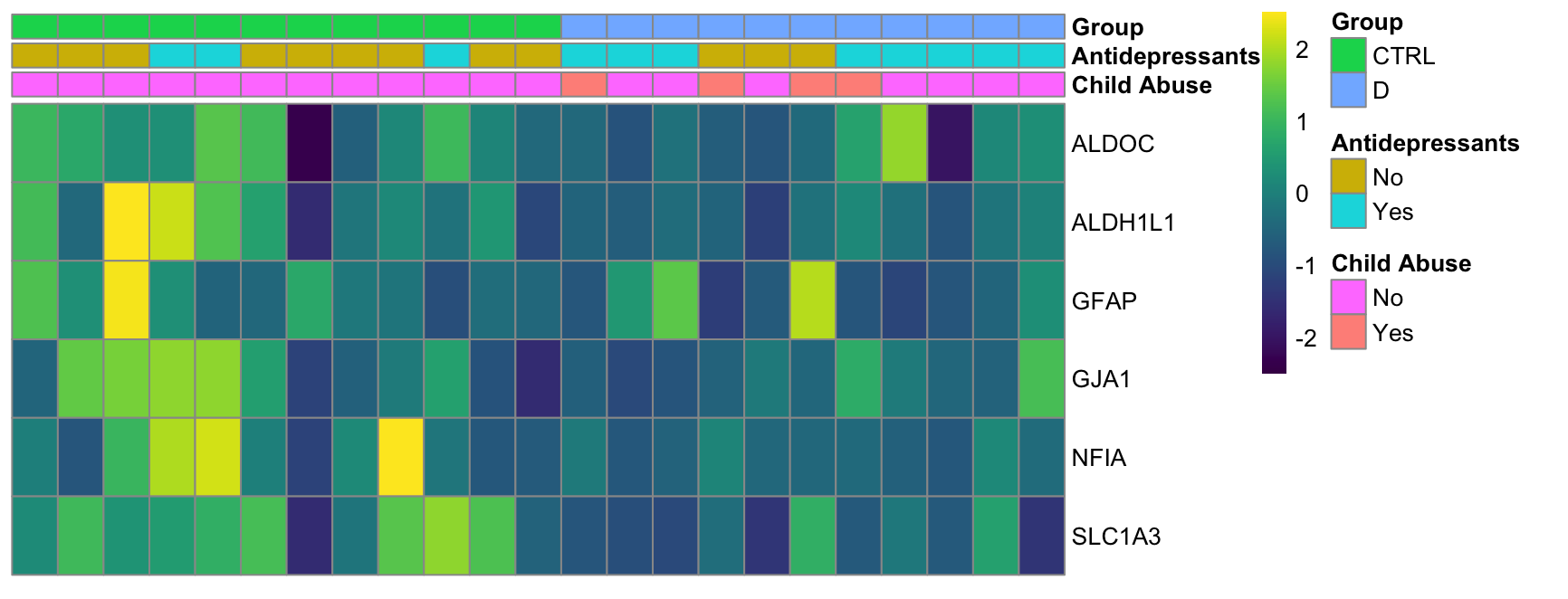

**Supplementary Table 1 – Human proteomics cohort**

|  | CTRL | D |
| --- | --- | --- |
| N | 12 | 11 |
| Axis 1 diagnosis | 0 | MDD (9); DD-NOS (2) |
| Age (years) (*P* = 0.27) | 68.25 ± 15.02 | 62.46 ± 8.82 |
| PMI (h) (*P* =  0.0004) | 36.07 ± 19.86 | 64.63 ± 9.70 |
| Tissue pH (*P* = 0.87) | 6.19 ± 0.30 | 6.21 ± 0.22 |
| Substance dependence | 0 | 4 |
| Antidepressants recently prescribed | 3 | 8 |

Data represent mean ± standard deviation. *P*-values were generated using unpaired *t*-test. D: individuals with depression; CTRL: healthy controls; DD-NOS: depressive disorder not otherwise specified; D: individuals with depression; PMI: post-mortem interval. Please note that the antidepressants in the control group were taken for non-MDD related conditions.

**Supplementary Table 2 - Phosphoproteomics cohort**

|  | CTRL | D |
| --- | --- | --- |
| N | 9 | 9 |
| Axis 1 diagnosis | 0 | MDD (7); DD-NOS (2) |
| Age (years) (*P* = 0.14) | 69.33 ± 16.75 | 60.11 ± 5.35 |
| PMI (h) (*P* = 0.0046) | 33.50 ± 21.80 | 61.84 ± 10.69 |
| Tissue pH (*P* = 0.85) | 6.16 ± 0.34 | 6.19 ± 0.30 |
| Substance dependence | 0 | 3 |
| Antidepressants recently prescribed | 3 | 7 |

Data represent mean ± standard deviation. *P*-values were generated using unpaired *t*-test. D: individuals with depression; CTRL: healthy controls; DD-NOS: depressive disorder not otherwise specified; D: individuals with depression; PMI: post-mortem interval. Please note that the antidepressants in the control group were taken for nonpsychiatric conditions.

**Supplementary Table 3 – FISH cohort**

|  | CTRL | D |
| --- | --- | --- |
| N | 3 | 5 |
| Axis 1 diagnosis | 0 | MDD (4); DD-NOS (1) |
| Age (years) (*P* = 0.59) | 48.0 ± 9.85 | 52.4± 11.80 |
| PMI (h) (*P* = 0.037) | 50.33 ± 7.29 | 76.46 ± 19.25 |
| Tissue pH (*P* = 0.89) | 6.16± 0.40 | 6.13± 0.26 |
| Substance dependence | 0 | 2 |
| Antidepressants recently prescribed | 0 | 2 |

Data represent mean ± standard deviation. *P*-values were generated using unpaired *t*-test. D: individuals with depression; CTRL: healthy controls; DD-NOS: depressive disorder not otherwise specified; D: individuals with depression; PMI: post-mortem interval.

**Supplementary Table 4 – IF cohort**

|  | CTRL | D |
| --- | --- | --- |
| N | 4 | 5 |
| Axis 1 diagnosis | 0 | MDD (3); DD-NOS (2) |
| Age (years) (*P* = 0.83) | 51.75 ± 11.00 | 52 ± 11.80 |
| PMI (h) (*P* = 0.064) | 47.75 ± 7.88 | 69.98 ± 14.18 |
| Tissue pH (*P* = 0.57) | 6.20 ± 0.34 | 6.22 ± 0.21 |
| Substance dependence | 0 | 2 |
| Antidepressants recently prescribed | 0 | 2 |

Data represent mean ± standard deviation. *P*-values were generated using unpaired *t*-test. D: individuals with depression; CTRL: healthy controls; DD-NOS: depressive disorder not otherwise specified; MDD: major depressive disorder; PMI: post-mortem interval.

**Supplementary Table 5 – Proteins detected before and after conservative cutoff**

| Dataset | # detected before cut-off | # detected after cut-off |
| --- | --- | --- |
| Human proteome | 4022 | 3702 |
| Human phosphoproteome | 7025 | 3957 |
| Mouse proteome | 3859 | 3844 |

**Supplementary Table 6 – Broad cell type markers for hypergeometric tests**

Provided as an excel file.

**Supplementary Table 7 – OB differentially expressed protein overlap with Maitra et al. (2023) differentially expressed astrocyte genes in males (snRNAseq)**

| Symbol | Name | Cluster name | dlPFC-RNA log2 D/CTRL | OB-Protein log2 D/CTRL |
| --- | --- | --- | --- | --- |
| AQP1 | Aquaporin-1 | Astrocyte 1 | -1.84 | -0.66 |
|  |  | Astrocyte 2 | -5.48 |  |
| CDC42EP4 | CDC42 Effector Protein 4 | Astrocyte 1 | -1.03 | -2.16 |
| MT1X | Metallothionein-1X | Astrocyte 1 | -1.21 | -1.32 |
| PLPP3 | Phospholipid phosphatase 3 | Astrocyte 1 | -1.13 | -0.72 |
| S100A1 | Protein S100-A1 | Astrocyte 1 | -1.08 | -0.76 |
| S1PR1 | Sphingosine 1-phosphate receptor 1 | Astrocyte 1 | -1.51 | -1.84 |
| SLC1A2 | Solute Carrier Family 1 Member 2 | Astrocyte 2 | -1.25 | -0.54 |
| SLC6A11 | Sodium- and chloride-dependent GABA transporter 3 | Astrocyte 1 | -1.08 | -0.72 |

**Supplementary Table 8 – Characterization of overlap in differential expression of human proteome and phosphoproteome**

|  |  | **Proteome** | **Phospho-proteome** | |
| --- | --- | --- | --- | --- |
| **Symbol** | **Name** | **log2 Abundance Ratio D/CTRL** | **log2 Abundance Ratio D/CTRL** | **Modification** |
| AHSG | Alpha-2-HS-glycoprotein | 1.11 | 2.31 | 1xCarbamidomethyl [C1]; 1xPhospho [S7(100)] |
| AQP1 | Aquaporin-1 | -0.66 | -3.83 | 1xPhospho [S19(100)] |
| DSP | Desmoplakin | 0.98 | 3.18 | 1xPhospho [S3(100)] |
| GFAP | Glial fibrillary acidic protein | -0.63 | -2.12 | 1xPhospho [S8(100)] |
| NEFH | Neurofilament heavy polypeptide | -0.64 | -2.13 | 1xPhospho [S6(100)] |
| SLC4A1 | Solute carrier family 4 member 1 | 1.45 | 3.06 | 1xPhospho [S10(100)] |
|  |  |  | 3.44 | 1xPhospho [S11(99.6)] |
| SPTA1 | Spectrin alpha chain, erythrocytic 1 | 0.86 | 2.33 | 1xPhospho [S3(100)] |
| THRAP3 | Thyroid Hormone Receptor Associated Protein 3 | -2.07 | -0.85 | 1xPhospho [S6(98.4)] |

**Supplementary Table 9 - Characterization of overlap in differential expression of human and mouse proteome**

|  |  | Human OB | Mouse OB |
| --- | --- | --- | --- |
| Symbol | **Name** | **log2 Abundance Ratio D/CTRL** | **log2 Abundance Ratio SD/CTRL** |
| AHSG | Alpha-2-HS-glycoprotein | 1.11 | 0.51 |
| ALB | Albumin | 0.73 | 0.57 |
| APOA1 | Apolipoprotein A-I | 1.34 | 1.48 |
| CAT | Catalase | 0.50 | 0.42 |
| CPT2 | Carnitine O-palmitoyltransferase 2, mitochondrial | -0.57 | -0.13 |
| CTNNB1 | Catenin beta-1 | -0.66 | -0.26 |
| DTNA | Dystrobrevin alpha | -0.87 | -0.27 |
| GHITM | Growth hormone-inducible transmembrane protein | 2.47 | 0.41 |
| HPX | Hemopexin | 0.59 | 1.09 |
| PSMD11 | 26S proteasome non-ATPase regulatory subunit 11 | 0.95 | -0.10 |
| TF | Serotransferrin | 0.80 | 0.37 |
| TIMM9 | Mitochondrial import inner membrane translocase subunit Tim9 | 2.30 | -0.24 |
| TTR | Transthyretin | 1.32 | 0.26 |

**Supplementary Table 10– Human OB proteomics processed spreadsheet**

Provided as CSV file

**Supplementary Table 11 – Human OB phosphoproteomics processed spreadsheet**

Provided as CSV file

**Supplementary Table 12 – Mouse OB proteomics processed spreadsheet**

Provided as CSV file
